# Zebrabase: An intuitive tracking solution for aquatic model organisms

**DOI:** 10.1101/248666

**Authors:** Jana Oltova, Jindrich Jindrich, Ctibor Skuta, Ondrej Svoboda, Olga Machonova, Petr Bartunek

## Abstract

Small fish species, like zebrafish or medaka, are constantly gaining popularity in basic research and disease modeling as a useful alternative to rodent model organisms. However, the tracking options for fish within a facility are rather limited. Here, we present an aquatic species tracking database, Zebrabase, developed in our zebrafish research and breeding facility that represents a practical and scalable solution and an intuitive platform for scientists, fish managers and caretakers, in both small and large facilities. Zebrabase is a scalable, crossplatform fish tracking database developed especially for research fish facilities but can be easily adapted for a wide variety of aquatic model organisms housed in tanks. It provides sophisticated tracking, reporting, and management functions that help keep the animal-related records well-organized including a QR code functionality for tank labeling. The implementation of various user roles ensures a functional hierarchy and customized access to specific functions and data. In addition, Zebrabase enables easy personalization of rooms and racks, and its advanced statistics and reporting options make it an excellent tool for creating periodic reports of animal usage and productivity. The communication between the facility and researchers can be streamlined by the requesting capabilities of the database. Finally, Zebrabase also features an interactive breeding history and a smart interface with advanced visualizations and intuitive color coding allowing to speed up the processes.

## Introduction

Zebrafish is a popular vertebrate model organism widely used in scientific research and a highly prized disease model. Its high fecundity, optical transparency, and rapid development make it very attractive for live imaging and developmental studies. Especially with the recent rapid progress in the field of genome editing, it seems to be reasonable to implement a database system that would facilitate tracking of the fish of F0 and F1 generations of the newly created, genetically modified lines, which can be both very laborious and space demanding and requires a high degree of organization in the facility. Also, proper outcrossing is required in zebrafish to prevent the inbreeding depression of the fish stocks^1^, which is a process particularly prone to mix-ups.

Despite the increasing abundance of the zebrafish model in research, tracking options suitable for the aquatic animals housed in tanks remain rather restricted, and scientists are often dependent on outdated software solutions or simple spreadsheets. One reason for that might be that the requirements for aquatic organism tracking differ quite significantly from the rodent tracking, where systems such as PyRat (https://www.scionics.com/pyrat.html) or open-source JAX Colony Management System^2^ (http://colonymanagement.jax.org) have long been established as a standard. The main particularity of aquatic animal tracking is group tracking, as opposed to the tracking of the individual animals. One of the reasons for group tracking is that up to date, there is no cost-effective system designed to distinguish single fish individuals in a tank. Technologies like subdermal chipping are developing fast, but the technology is not yet miniaturized enough to enable routine use in small fish species, although some significant advances have been reported using the RFID microtags^3^. Also, because of the social requirements and bulk mating, zebrafish are recommended to be kept within a specific density range in the tanks with both males and females present ^4,5^. Keeping the individual fish in solitude has a negative impact on their development and health ^6^ and is, moreover, very space inefficient.

There is a handful of existing solutions, either commercial ones (e.g., http://www.daniodata.com/, https://www.scionics.com/pyrat_aquatic.html) or opensource^7-9^ http://zebase.bio.purdue.edu/, http://aqacs.uoregon.edu/zf/files/, https://zebrafish.jimdo.com/), but none of these seems to be prevalent in the zebrafish community, probably because each of them suits best to a slightly different set of requirements of the facilities. Two main drawbacks of the solutions currently in the market are the price and dedication to a single platform, which can both be perceived as a limiting factor, especially for small facilities with a limited budget. The sustainability and implementation of new functions have proved as the bottleneck of some of the databases that were developed in the past years and indeed, this aspect should be considered carefully, as switching from one system to another can be quite tedious, especially if the tracking history has to be kept. Lastly, as the reporting requirements from local authorities are getting more rigorous in many countries, another feature which is missing in some of the existing solutions is a detailed fish usage and health reporting capability.

## Results

### Basic operation principles

Zebrabase enables to keep track of the animals from birth to death and store detailed records of all the relevant events. Every action performed by the user creates a record in the database and can be later used for reporting purposes. Zebrabase is optimized for touch devices and supports the use of QR codes for tank labeling to enable efficient tracking of the animals.

### Fish characterization and grouping

The most important purpose of Zebrabase is the time-efficient and comprehensive tracking of the fish stocks. Therefore, all the animals in the facility are divided into fishlines, which are specified by the database administrator by defining its genotypes, the date of creation or import and other optional information (https://public.zebrabase.org/fishline/create). Fish corresponding to a single fishline are further organized into groups of siblings, stocks, sharing the same date of birth and parents. As in some stocks the number of fish might exceed the capacity of a single tank, we have introduced another level of grouping animals, substocks, which refer to a defined stock subset in a single tank. Thus, substock is the basic unit that is tracked in the database and can consist of a variable number of fish individuals.

Genotypes consist of modification category (transgene, natural mutant, engineered mutant or custom) and either driver, triggered gene and an optional allele for transgenic genotypes or affected gene and allele/mutation for mutant genotypes. With slight modifications required by the automatic zygosity calculation algorithm, the Zebrabase name generation rules are based on the common ZFIN nomenclature (https://wiki.zfin.org/display/general/ZFIN+Zebrafish+Nomenclature+Guidelines). It is also possible to specify an alias, which can be used as a trivial name for substocks with very long generated names (e.g., in the case of multiple genotypes), or a suffix appended to the generated name.

Zebrabase enables to store attachments for each of the fishlines to effectively organize genotyping protocols, publications, and visual content associated with it. These can be for example phenotypes of mutant lines or transgene expression patterns that can serve as a sorting reference for reporter lines. To achieve an additional degree of organization of the fishlines, responsible user, and a workgroup can be specified.

### Spatial and temporal tracking

We have developed a system for tracking animals in the facility, which utilizes the positional widget called Facility (Fig. 1A), a platform for user-defined rooms, racks, and tanks. In Facility, the productivity of the substocks is visualized by consistent color coding to provide a quick status reference for the users. We have also optimized the QR code functionality to record the action history efficiently for each of the substocks via any mobile device, as an alternative to the manual entering from a desktop computer. In addition, a list view of all the substocks regardless of their position in the facility, including deceased or terminated substocks, can be found under the Fish tab (Fig. 1B).

**Figure 1.**
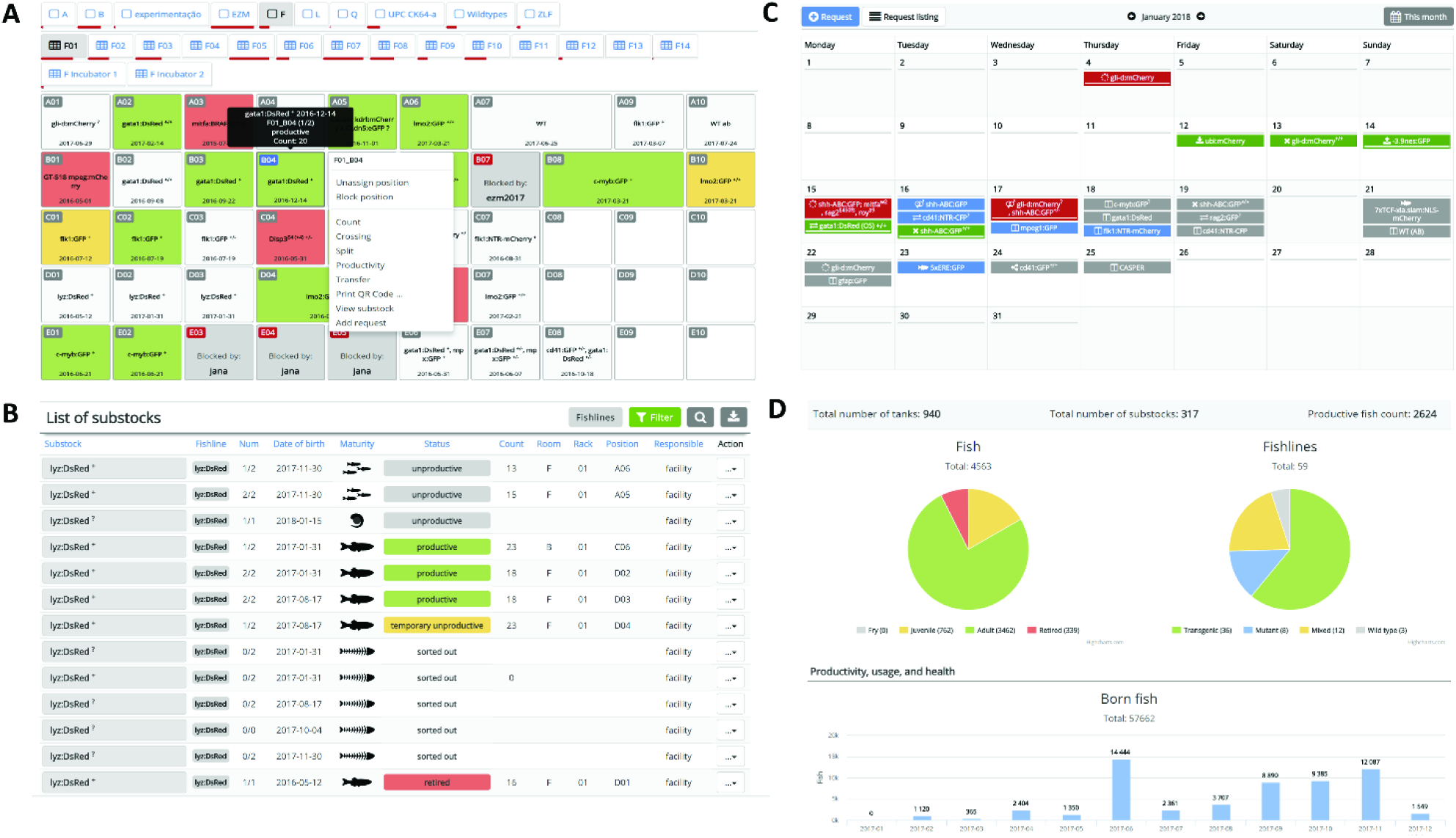
Zebrabase user interface. **A)** Facility view is a mobile-optimized visual facility representation that enables to browse all the positions within the facility and perform all the actions. Rows are designated by letters and horizontal positions by numbers. Combined with the rack ID, each position is uniquely defined within the whole facility. Upon click, the action menu appears that enables to perform all the actions available for the particular substock. **B)** Fish list is a list of all the records in the facility regardless of whether they have a specific position in the facility or not. Records that can be intuitively filtered according to various parameters to create a subset of fish of our interest. By default, only active (alive) fish are shown in the list, but that can be changed by adjusting the filtering options. Substocks are color-coded in the following manner: green – productive, dark grey - blocked, light grey – unproductive, red - retired, yellow - temporary productivity issues. **C)** Representation of various requests and self-requests in the calendar. Various icons are used for various types of requests (fish, setup fish, crossing, terminate, genotyping, mobilize fish, transfer, import, export). The requests are color-coded in the following manner: grey – new requests, blue – accepted requests, green – performed requests, red – declined requests. **D)** Zebrabase statistics enables to store, visualize and export data for the reporting purposes. The overview of the facility statistics is presented in the form of pie charts. Left – overview of the age distribution of the fish in the facility. Right – overview of the fishlines in the facility. Bottom - the bar charts are provided and plotted monthly for fish born in the facility, deceased fish, euthanized fish and fish used for experiments (images not shown).

An interactive breeding history diagram is shown at the substock record page https://public.zebrabase.org/substock/1673), which provides links to all the substocks in the whole pedigree. It also shows the dates of crosses and splits and, if clicked, leads to the action detail page. Breeding history enables to efficiently track animals that potentially shared the same housing or breeding tank in the past (e.g. during outcrosses), when a cautious approach needs to be adopted upon disease outbreak, potential mix-ups of fish, etc.

Zebrabase also features a number of actions – Crossing, Split, Transfer, Count, Productivity, Genotyping, Add Request and Print QR code – that are fully trackable and can be browsed or filtered at any time in case a specific event needs to be traced back. Action history is also listed on the substock record page to provide a full overview of the actions that have been performed with a particular substock.

### Age of the animals and productivity

We have implemented statuses for each substock that are determined by productivity (the ability to spawn), age and additional information inserted by the facility staff or researchers. Each of these statuses is visualized in a specific color to highlight the problematic substocks so that they are not used in an experiment by mistake. These basic statuses are Unproductive, Productive, Retired, Deceased, and Sorted out. Moreover, there are temporary statuses that can be turned on and off to visualize temporary productivity or progeny survival issues. The specific reason for the temporary statuses should be defined by each facility independently based on their operating standards.

### Breeding management

The Crossing action serves for incrossing and outcrossing of the substocks in the database. When outcrossing substocks of different fishlines, new fishline with corresponding genotype combination is created automatically, unless it is already present in the database. In the same manner as all the other actions, Crossing can be initialized both from the Fish list or the Facility view. Moreover, Zebrabase enables to distinguish automatically whether fish are hetero-or homozygous or unknown zygosity for a specific genotype based on the zygosities of the parents. To plan or request crosses, the requesting functionality is accessible via the Calendar tab.

### Genotyping

For the purposes of genotyping and sorting of the fish according to the transgene or mutation presence, there are two functionalities that help to improve the workflow. The action Genotyping enables to enter zygosity independently for each of the genotypes. If a more complex, physical rearrangement of the fish substocks is performed, a compound action Split enables to split the fish stock according to the genotypes, zygosities or any other parameter and at the same time, enter their counts and positions.

### Workgroups and user rights

In case more research groups are sharing the facility and wish to have their data separated, the workgroup functionality enables to setup multiple workgroups with defined administrators, whose data will be completely separated in the user interface although they are using a single database instance. However, the facility members (e.g., caretakers) can still retain the right to see the data of all workgroups sharing the facility, if required. Within each workgroup, there are various user roles that ensure different rights assigned to the user. Administrator, facility staff, standard user and guest rights with different permissions from view-only to the most advanced administrator rights can be assigned to every user, where a single user can have more user right groups assigned, and the permissions then combine.

### The calendar, requesting and experiment planning

Submitting a request is a way to ask the facility staff for specific actions like setting up the fish for crossing, splitting or terminating substocks and fishlines. Every such a request is then recorded in the database and becomes visible in the Calendar (Fig. 1C). Email notifications for new requests can be sent to the facility staff emails, if configured. The facility members are then able to accept the request, mark it as performed when the request is completed or decline it. Using the same functionality but assigning self-requests is a useful way to plan the experiments. Self-requests are visible only to the user who created it. Generally, there are several request types that can be entered: Fish, Setup Fish, Transfer substock, Terminate substock, Import, Export, Crossing, Mobilize Fish, and Genotyping.

### Statistics and reporting counts and usage of the animals

Zebrabase utilizes a comprehensive system of reporting animal usage or death via the action Count. At first, the start count needs to be specified, which is typically performed when the juvenile fish are introduced into the facility system, but can also be done any time earlier or later. Once this figure is available for a particular substock, it is possible to subtract used or deceased fish and specify for which reason they have been sacrificed, if appropriate (e.g., type of an experiment). These records are then visualized in the Facility overview (Fig. 1D) and can be used to generate reports.

### Data import and export

We provide a simple spreadsheet template that enables to import a complete dataset including the breeding history to a new Zebrabase instance. Data exporting options include the export of statistics and the full substock data into an ․xls or ․csv files and upon request, complete database dumps can be provided.

## Methods

### System architecture

Zebrabase is a web-based, cross-platform hosted zebrafish tracking solution. Zebrabase supports both desktop and mobile use on Mac, Linux, Windows, Android and iOS and has a built-in QR code generator and QR code reader functionality that provide instant access to all the fish stock information and related actions on a mobile device. The Zebrabase system is operated with the utmost effort to ensure service availability. The system is running on a virtual server in the environment composed of two clusters of hardware servers. Individual hardware components of the environment are redundant, and each cluster is housed in a different data center of the Institute of Molecular Genetics. The system is replicated from the primary to the secondary cluster every 30 minutes. In case the primary cluster becomes unavailable, the service can be restored in a short time on the secondary cluster by a system administrator.

### Database development

For every instance of Zebrabase, a dedicated PostgreSQL database https://www.postgresql.org/) is created to ensure high data security and consistency. The code base (Zebrabase version), both back- and front-end, is the same for all Zebrabase instances. For the back-end part of the application, the Python programming language https://www.python.org/) is employed with the Django web framework https://www.djangoproject.com/) that facilitates the data exchange between a web browser and the database. The front-end part is implemented using latest HTML5/CSS3 technologies that enables the interface to be responsive, interactive and to make use of data visualizations. Since Zebrabase is a web application, it is multi-platform (Windows, Mac OS, Android, iOS, Linux-based systems) and works in all modern internet browsers (Google Chrome, Mozilla Firefox, Microsoft Edge, Apple Safari, etc.).

All communication of the users with the Zebrabase system is secured via SSL, and for each customer (e.g., facility or laboratory), a completely separate instance of Zebrabase is provided. Database dumps are performed four times a day and archived for 6 months, in the case that data restoration is necessary. Additionally, the fish stock records and statistics can be exported to an .xls or a .csv file directly from the user interface at any time to provide additional data backup.

### User interface

Zebrabase user interface provides full touch support and the cross-platform access via web browser. Zebrabase implements an intuitive and consistent color coding, icons and visualization in the GUI that enables easy orientation in the fish substocks. The user support is reachable via the Feedback link at the bottom of every Zebrabase instance, which also provides an interface for suggestions of new features or improvements.

## Discussion

As the popularity of small fish species in research is rising, the need for a proper tracking database that would allow to keep records of all the animals, provide statistics and reporting capabilities, and enhance communication of the user and the caretakers in larger scale breeding facilities is critical. In this article we propose a novel tracking database for aquatic model facilities, especially optimized for the use with small fish species like zebrafish. There were several major shortcomings of the existing solutions that led us to the idea of the development of Zebrabase, like commitment to a single platform, limited data export and transfer options, poor sustainability, or missing mobile device support. With Zebrabase, we tried to tackle these issues and create a tracking solution that will suit the needs of aquatic model facilities of any scale.

Zebrabase has been built around several key concepts that guarantee cross-platform operation including full mobile device support and hosted, maintenance-free service for the users, unparalleled by any other solution. The main cornerstones of Zebrabase are comprehensive animal tracking, interactive breeding history, and advanced management features. The animal tracking system of Zebrabase has been designed with attention to the crucial details that make the zebrafish tracking different from the tracking of other model organisms - above all, following the group of animals as opposed to individual animal tracking, which determines substantially the way the data are handled and interpreted.

Next, we put an emphasis on the reporting capabilities of the database, which allow to store fish census and usage reports for the whole facility in a single Excel file. This report provides a monthly overview of all the fish born, deceased, euthanized and used for experiments. We have also reflected the suggestions of our beta testers and enhanced the requesting and experiment planning features of the database. The latest version now enables to assign various request types either to the facility or user himself, so that all the actions that need to be performed in the fish facility are comprehensibly visualized. The advanced management features of Zebrabase, like workgroups, user roles, responsible users, and QR code labeling will be a great benefit, especially for larger sized facilities. Full touch and camera integration and QR code support are provided in the recent Zebrabase version, to ensure maximum versatility of the system and accessibility of the data from any device, which can be very beneficial for the real-life use in the fish facilities. In-App camera support is currently unavailable on Apple devices but the substock records can be alternatively accessed via any mobile QR code reader application.

Zebrabase is a non-profit project designed to support small and starting facilities by providing the service for free for up to 150 active substocks or 3 racks. After reaching this threshold a yearly fee covering the initial setup, data backup and recovery, version updates, server maintenance and user support is charged, which corresponds to the size of the facility. For details on pricing please visit the Zebrabase webpage (https://zebrabase.org/faq). To obtain the demo access or request a new database instance, please use the web form at https://zebrabase.org/contact.

Thanks to the hosted character of the Zebrabase project, we are able to provide unparalleled possibilities, like automatic backup and recovery options or instant version upgrades that are automatically available for all the instances, regardless of whether they are running on the free or paid plan. The project sustainability is guaranteed thanks to the fact that the tool is in use in the fish facility of the Institute of Molecular Genetics, where most of the new features are designed and thoroughly tested. The feedback from the users is regularly collected to help the users decide which new features should get the highest priority and the users are encouraged to report new suggestions for improvement to the Zebrabase team. In the near future, we plan to implement batch actions, enterprise resource planning and genome editing module to streamline the workflow of researchers and facility managers and enable charging the users for the facility service. For more information about the new functionalities, please, visit our webpage https://zebrabase.org/news).

## Acknowledgement

This work was supported by the Czech Science Foundation (16-21024S) and the Ministry of Education, Youth and Sports (NPU I – LO1419).

